# Treatment of *Saccharomyces cerevisiae* with cigarette smoke extract causes vacuolar fragmentation to combat cigarette smoke-induced cellular toxicity

**DOI:** 10.64898/2026.01.30.702977

**Authors:** Aditya Shukla, Srimonti Sarkar, Alok Kumar Sil

## Abstract

Exposure to cigarette smoke is one of the major risk factors for developing various diseases such as chronic obstructive pulmonary disease (COPD), cardiovascular disorders, and cancer mediated via cellular oxidative stress and organelle dysfunction. To this end, the current study investigated how cigarette smoke extract (CSE) affects vacuole structure and function in *Saccharomyces cerevisiae*, as vacuole plays a crucial role in handling oxidative stress–induced misfolded proteins. Our results showed that CSE exposure causes transient vacuolar fragmentation up to 1 h to increase its surface area to facilitate microautophagy in clearing CSE-mediated misfolded protein and promoting cell survival. However, excessive fragmentation or vacuolar fusion sensitizes cells towards CSE-mediated cellular toxicity. Towards understanding the underlying mechanism, the current study demonstrated the involvement of PI3P and PI (3,5) P_₂_-mediated signaling and phospholipase-driven remodeling of lipid moieties. Moreover, the current study also showed the importance of mitochondrial activity in CSE-mediated vacuolar fragmentation. Prolonged exposure to CSE impairs mitochondrial function and thus disrupts fragmentation, the adaptive survival strategy against CS. It results in proteostasis collapse, which is a characteristic shared by many inflammatory and degenerative disorders. Taken together, the current study reveals a previously unrecognized cellular protection mechanism induced by cigarette smoke and highlights potential therapeutic targets for mitigating CS-mediated diseases

## 1. Introduction

Cigarette smoke (CS) is a mixture made up of around 4,700 different chemicals, such as heavy metals, aldehydes, a variety of reactive oxygen and nitrogen species (ROS and RNS), and several polycyclic aromatic hydrocarbons. These oxidants include relatively stable organic radicals like semiquinone and short-lived forms like superoxide anions [1]. These substances damage proteins, lipids, and nucleic acids, which either directly or indirectly causes oxidative stress. In addition to its contents, CS can induce the generation of various oxidants at the cellular level. It reacts chemically with biomolecules to produce more oxidants, activates endogenous ROS-producing systems, and concurrently compromises antioxidant defenses [2]. Thus, in contrast to single-agent oxidants like hydrogen peroxide (H2O_₂_), CS-mediated oxidative stress is more robust and due to its multidimensional action, it overwhelms cellular redox equilibrium. Therefore, exposure to CS is associated with significant cellular damage and an increased risk of conditions such as atherosclerosis, cancer, and chronic obstructive pulmonary disease (COPD) [3,4].

Oxidative stress and proteotoxicity induce adaptive remodeling of cellular compartments at the organelle level by altering basic homeostatic processes. Numerous organelles undergo membrane fission and fusion events during cell division, vesicular traffic, or in response to environmental changes. A few examples are the lysosome [5], peroxisomes [6], mitochondria [7], and golgi [8]. The morphology of vacuoles dynamically adjusts to intra- and extracellular environmental conditions by fission, also known as fragmentation, and fusion, which results in the formation of either multiple or a single, larger vacuole respectively. For instance, in *Saccharomyces cerevisiae*, vacuole fragmentation is caused by lactic acid stress, endoplasmic reticulum stress, or hyperosmotic shock, whereas vacuole fusion is induced by hypotonic shock, nutrient limitation, or TORC1 inactivation [9, 10]. Vacuolar fragmentation also occurs during cell division, so that vacuoles can be partitioned into daughter cells [11].

In *Saccharomyces cerevisiae* vacuoles have roles in osmoregulation, autophagy, as well as in the storage of various ions and amino acids [12]. Lysosomal disorders are caused by mutations in genes that affect lysosomal degradation, indicating the importance of regulating lysosomal/vacuolar hydrolytic capacity to the proper functioning of eukaryotic cells [13].Previous reports have shown oxidative stress causes accumulation of misfolded proteins and under this stressed condition, vacuolar/lysosomal function (autophagy) in tandem with the ubiquitin-proteasome system is vital to maintain proteostasis [14]. In this context, recent report has shown that enhanced microautophagic activity is upon vacuolar fragmentation, however, it does not influence the macroautophagic activity [15]. Therefore, fragmentation, which leads to an increase in the surface area of vacuolar membranes, might play an important role in the clearance of accumulated misfolded proteins. This could be another cellular mechanism to eliminate the toxic effects resulted from accumulation of misfolded proteins. . Since CSE-mediated oxidative stress promotes protein misfolding and many of the CSE-mediated diseases are caused due to an accumulation of misfolded proteins the current study investigated the effect of cigarette smoke induced-oxidative stress on yeast vacuolar structure and its function. The results showed that the cell treated with CSE undergoes fragmentation at the beginning of the treatment, and it contributes to better clearance of misfolded proteins and better survival. However, results also showed that too much fragmentation is not good for cellular survival.

## 2. Results

### 2.1. Treatment of *S cerevisiae* cells with cigarette smoke extract (CSE) induces vacuolar fragmentation

To investigate the effect of cigarette smoke on vacuolar physiology, yeast cells were treated with CSE for different periods as indicated in the figure and stained the vacuoles with FM4-64 dye, which is widely used to stain vacuolar membrane [20]. After CSE treatment, cells exhibited fragmentation of vacuole at 0.5 h and thereafter it is reduced progressively with time (Figure 1A, Left Panel).

**Figure 1.**
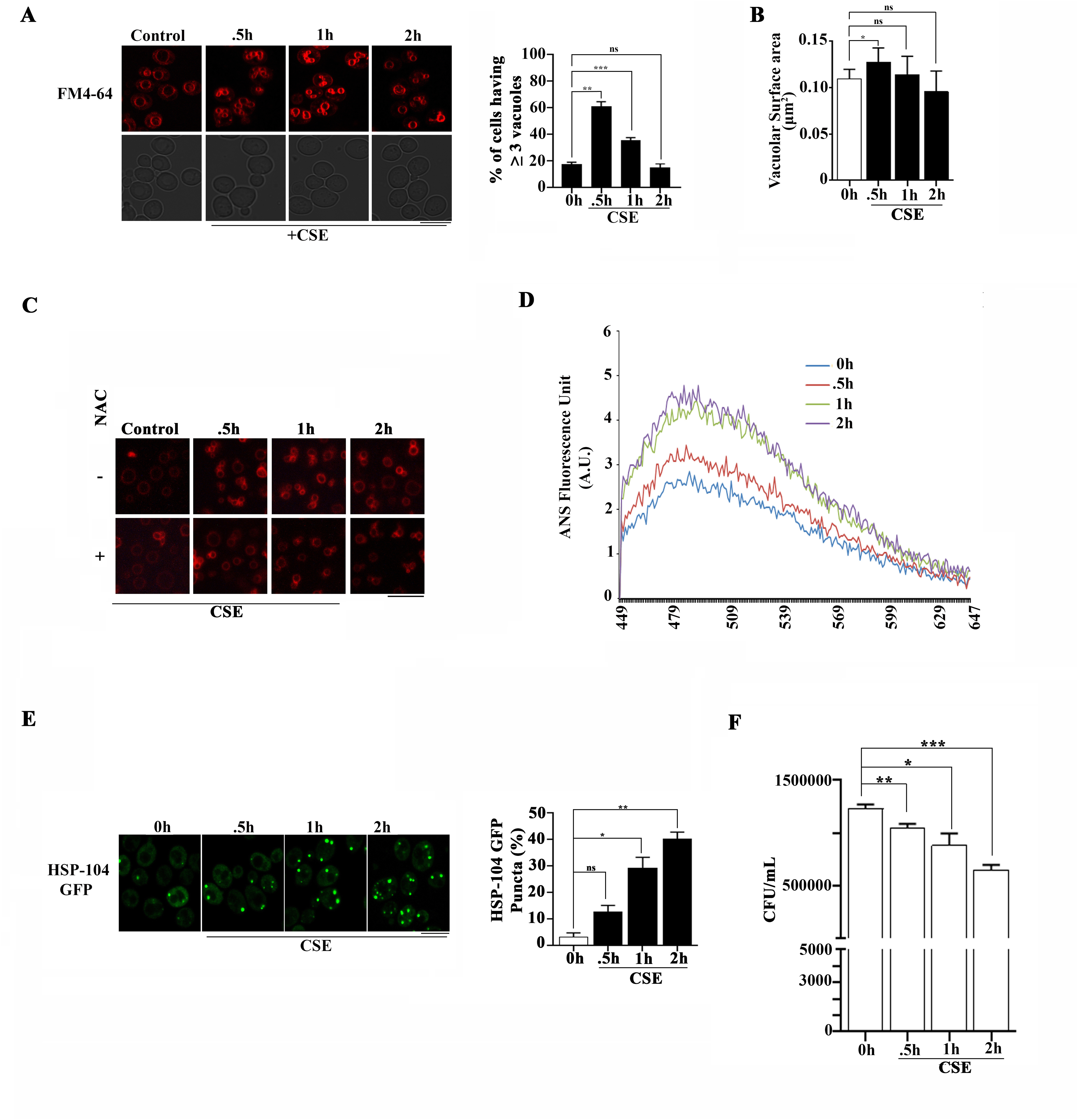
CSE-induced oxidative stress causes vacuolar fragmentation: A) CSE induced vacuole fragmentation profile. Cells were grown till log phase and were then stained with FM4-64 for 30 mins followed by chasing for 1 h in YPD media. Cells were then treated with CSE for different time periods as indicated and visualized under microscope. Vacuolar morphology of approximately 150 cells was calculated. The bar diagrams at the right represents the quantitative profile. B) Vacuolar surface area profile. after CSE treatment. Surface area of vacuoles was calculated using Imagej software (NIH).For each condition, approximately 10 different fields containing atleast 30 cells for each field. C) NAC prevents CSE-induced vacuolar fragmentation. Cells were grown till log phase and were then stained with FM4-64 for 30 mins followed by chasing for 1 h in YPD media. Cells were then differentially treated with CSE and 10mM NAC for different time periods as indicated and visualized under fluorescence microscope. D) Misfolded protein profile by ANS staining. Cells were grown until the early exponential phase and adjusted to an OD_600_ of 0.1. Cells were treated with 15% CSE for different time periods. Cells were then stained with ANS (30μM) dye and the level of misfolded protein was determined spectrofluorimetrically. E) Misfolded protein profile with HSP-104 GFP. Cells harboring HSP104-GFP were grown till mid log phase and then treated with CSE for different time period, and visualized under confocal microscope The bar diagram at the right represent quantitative profile. F) Cell viability profile. Cells were grown until the early exponential phase and adjusted to an OD_600_ of 0.1, followed by 15% CSE treatment for different time period as indicated. In all the cases, Serial dilutions were spotted on YPD plates, following which the plates were incubated at 30 °C for the indicated time periods (48 or 72 h) before scoring for growth. Each image is representative of ten different fields. Scale bars: 8μm. Results are represented as mean ± SD of data obtained from 10 different fields using paired t test (n.s., not significant; ∗p value < 0.05; ∗∗p value < 0.005; ∗∗∗p value < 0.0005).. (a.u. indicates arbitrary unit).

Quantitative profile also supported this view (Figure 1A, Right Panel). The likelihood of fragmentation at the beginning of treatment is to increase the surface area, which may facilitate increased contact between the misfolded protein-harboring endosome and the vacuole, thereby promoting their removal from the cell and helping the cell to combat CSE-induced oxidative stress. To this end we measured the vacuolar surface area to volume ratio after CSE treatment and found that at 0.5 h the surface area of the vacuole was increased with respect to the control, and thereafter it decreased progressively with time (Figure 1B). To confirm further that CSE-mediated oxidative stress is the cause of this defect, cells were treated with N-acetylcysteine (NAC), a known antioxidant, in conjunction with CSE and then observed for vacuolar morphology. The presence of NAC antagonized the effect of CSE, and the morphology of vacuole was comparable to wild type control cells (Figure. 1C). To verify whether increased vacuolar surface area is correlated to better clearance of misfolded protein, we examined the amount of misfolded protein at different time points using ANS dye, ANS dye monitors misfolded proteins by binding to exposed hydrophobic regions that become more accessible when proteins unfold or misfold. This enhanced binding thus increases the fluorescence intensity and provides a useful tool for measuring misfolded proteins. The result showed that the amount of misfolded protein increased with time and it perfectly matches with the vacuolar fragmentation events within the cell (Figure 1D).To further verify accumulation of misfolded proteins after CSE treatment, yeast cells harboring HSP104-GFP gene was treated with CSE and thereafter examined the presence of bright puncta in the cytosol. Hsp104, a disaggregase protein, interacts with misfolded protein aggregates and facilitates either the refolding of these misfolded proteins or their degradation [21]. Thus, the presence of cytosolic puncta in the CSE-treated transformant can be used as a marker for aggregated misfolded proteins. Consistent with the ANS data, no or very few green puncta were observed in the control cells that were not treated with CSE. On the other hand, CSE-treated cells exhibited the presence of more green cytosolic puncta compared to control and the number was markedly increased after 1 and 2 h of treatment (Figure 1E, Left Panel). Quantitative profile also supported the view (Figure 1E, Right Panel). Since increased fragmentation upon CSE exposure contributes to better misfolded protein clearance, we speculated that cellular survival will also be positively correlated with vacuolar fragmentation. So cellular viability assay was performed, and congruent with our expectation, least cell death was observed for 30 mins and thereafter, cell viability progressively decreased with time (Figure 1F).

If vacuolar fragmentation helps to combat toxicity, then agents that promote vacuolar fragmentation should exhibit enhanced resistance against CSE-mediated toxicity. To verify this, we pre-treated cells with 0.5 mM NaCl before exposure to CSE, as NaCl is a known inducer of vacuolar fragmentation [22]. Consistent with the expectation, more vacuolar fragmentation was observed for cells treated with NaCl and CSE than for cells treated only with CSE (Figure 2A, Left Panel). Quantitative profile also supported this view (Figure 2A, Right Panel). Consistent with this, cells treated with both NaCl and CSE exhibited reduced accumulation of misfolded proteins than cells treated only with CSE and consequently a better survival against CSE toxicity was observed for cells treated both with CSE and NaCl than cells treated only with NaCl (Figure 2B, C).

**Figure 2.**
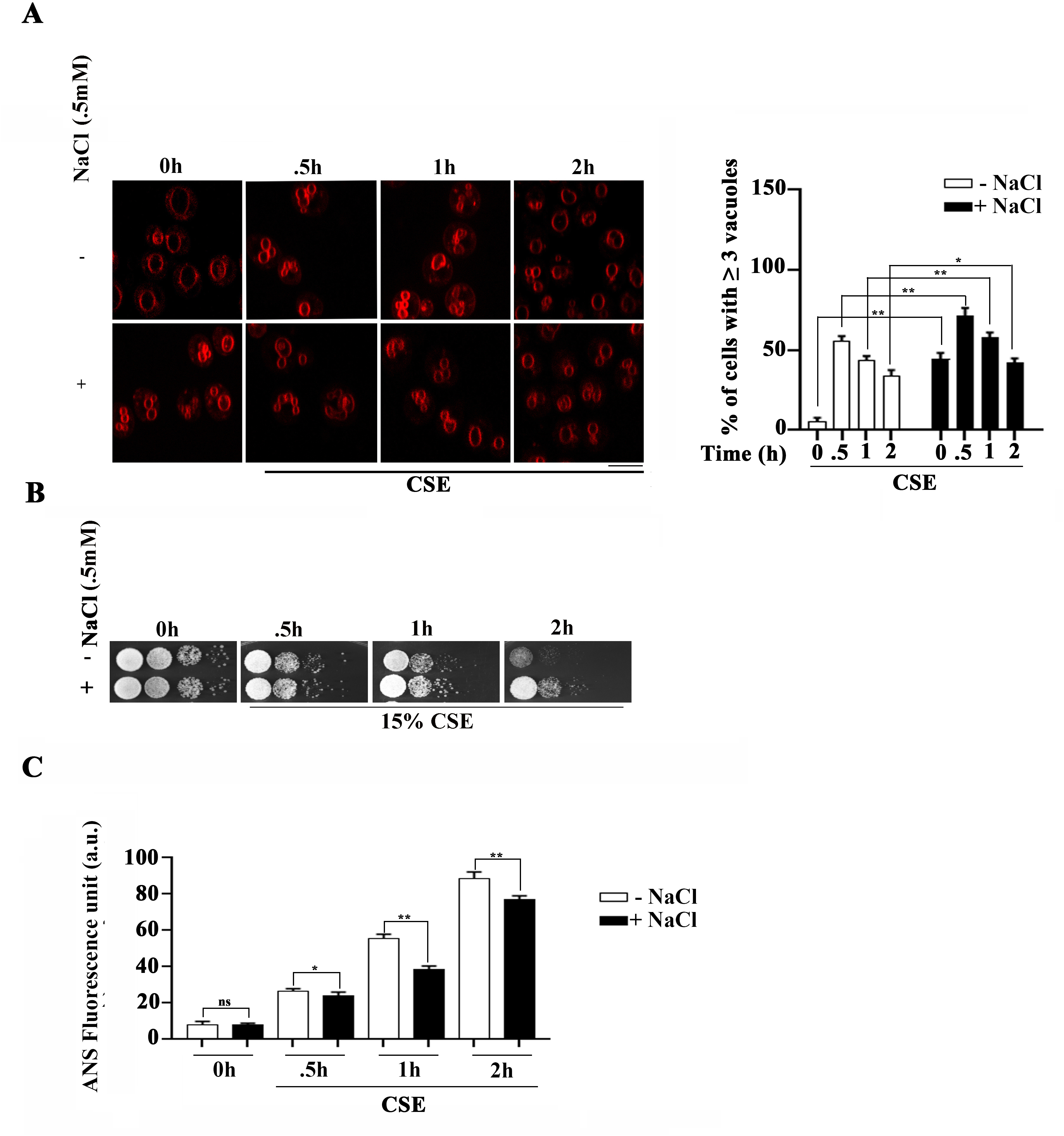
Vacuolar fragmentation helps combating CSE induced cellular toxicity. A) NaCl enhances vacuolar fragmentation. Cells were grown till log phase and were then stained with FM4-64 for 30 mins followed by chasing for 1 h in YPD media. Thereafter, cells were treated with 0.5 M NaCl for 15 mins followed by CSE treatment for different time period as indicated and visualized under microscope. Vacuolar morphology of approximately 150 cells was calculated. The bar diagram on the right side represents quantitative profile. B) Spot assay profile. Exponentially growing cells treated with CSE in the presence and absence of NaCl for different time intervals and were then spotted on YPD agar plates. C) Misfolded protein profile. Cells were grown until the early exponential phase and adjusted to an OD_600_ of 0.1. Cells were treated with 15% CSE for different time periods. Cells were then stained with ANS (30μM) dye and the level of misfolded protein was determined spectrofluorimetrically. Each image is representative of ten different fields. Scale bars: 8μm. Results are represented as mean ± SD of data obtained from 10 different fields using paired t test (n.s., not significant; ∗p value < 0.05; ∗∗p value < 0.005; ∗∗∗p value < 0.0005). (a.u. indicates arbitrary unit).

### 2.2. Vacuolar fusion makes the cell sensitive to CSE-induced toxicity

It is likely that if vacuolar fragmentation promotes cell survival against CSE-induced toxicity, vacuolar fusion will make the cells sensitive to CSE. To verify this, we examined the effects of vacuolar fusion on cell survival upon CSE treatment. For this purpose, yeast cells were transformed with *YPT7,* which is known to promote vacuolar fusion, and subsequently these transformant were treated with CSE for different periods. Vacuolar morphology, misfolded protein level, and cell survival were examined as before. Interestingly, Ypt7 over-expressing cells exhibited vacuolar fragmentation with cellular survival almost similar to WT cells at 0.5 h. Thus, despite the presence of Ypt7, cells promote vacuolar fission over fusion upon CSE-induced oxidative stress (Figure 3A, Left Panel). Quantitative profile also supported this view. (Figure 3A, Right Panel). Consistent with the vacuolar fragmentation, at 1 and 2 h, cellular survival was better for WT cells compared to *YPT*7overexpressing (Figure 3B). This result again underscores the importance of vacuolar fragmentation to combat CSE-mediated toxicity. Additionally, consistent with the vacuolar fragmentation the overexpression of *YPT7*, exhibited significantly higher levels of misfolded proteins compared to wild-type cells at later time points. These results indicate that CSE induces vacuolar fission that helps to combat CSE-induced cellular toxicity (Figure 3C).

**Figure 3.**
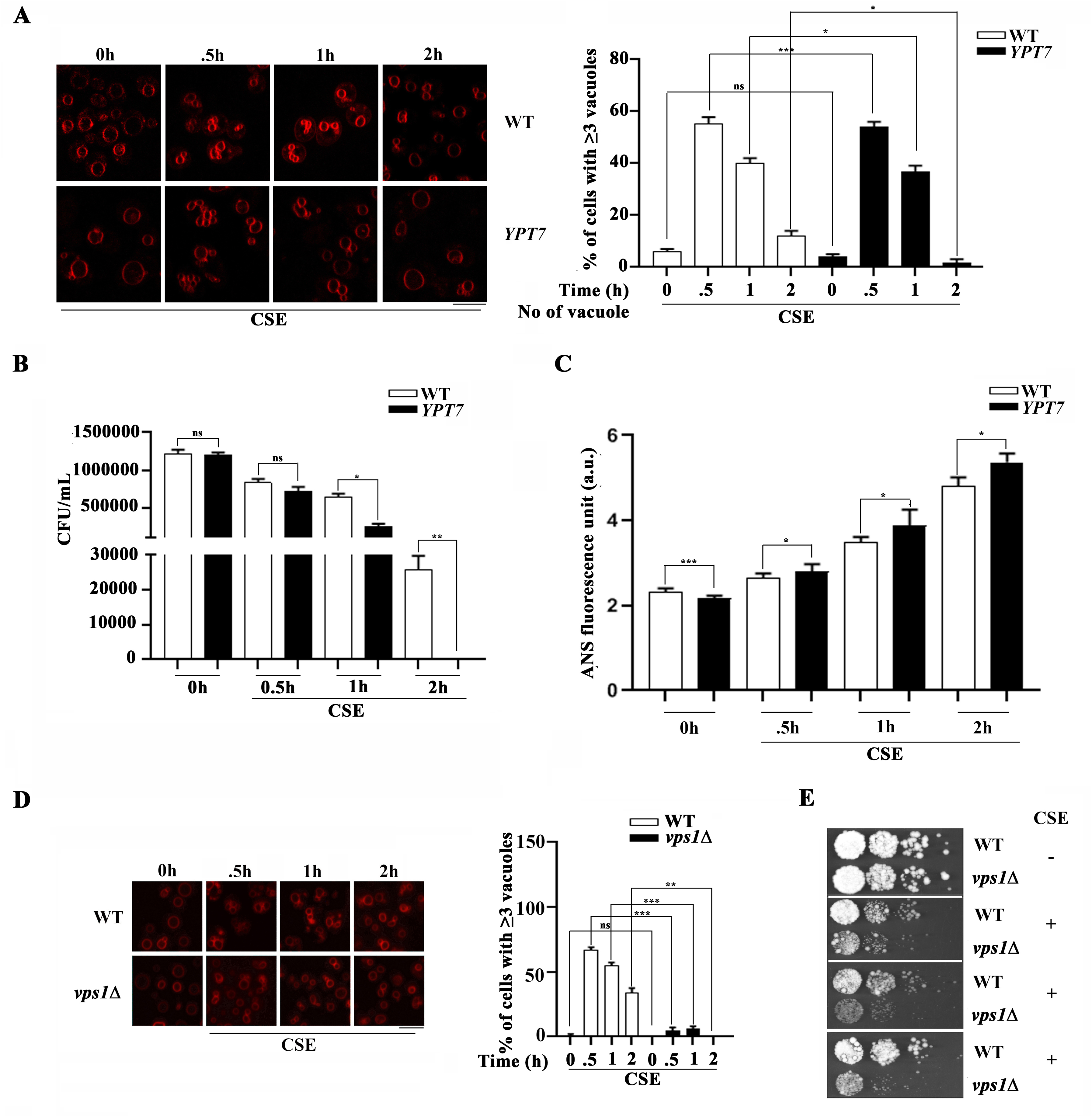
Vacuolar fusion sensitizes cells to CSE. A) CSE-induced vacuolar fragmentation profile in presence of Ypt7. Cells harboring or were not harboring *YPT7* were grown till log phase and then stained with FM4-64 for 30 mins followed by chasing for 1 h in YPD media, centrifuged and resuspended in YPD..Cells were then treated with CSE for different time period as indicated and visualized under microscope. Vacuolar morphology of approximately 150 cells was calculated. The bar diagrams at the right represents the quantitative profile. B) CFU profile. Exponentially growing cells harboring or were not harboring *YPT7* were treated with CSE at different time intervals, cells were then centrifuged and resuspended in YPD followed by CSE treatment. Serial dilutions were spotted on YPD plates, following which the plates were incubated at 30 °C for the indicated time period 72 h. C) Misfolded protein profile. Cells harboring or were not harboring *YPT7,* were pre grown at 30°C until the early exponential phase and adjusted to an OD_600_ of 0.1. Cells were then treated with 15% CSE for different time period as indicated and incubated with 30 μM ANS dye for 30 minutes. The level of misfolded protein was determined spectrofluorimetrically. D) Vacuolar fragmentation is Vps1 dependent. WT and *vps1*Δ cells were grown till log phase and were then stained with FM4-64 for 30 mins followed by chasing for 1 h in YPD media. Cells were then treated with CSE for different time period and then visualized under microscope. Vacuolar morphology of approximately 150 cells was calculated. The bar diagram at the right side represents quantitative profile. E) Spot assay for *vps1*Δ cells. Exponentially growing cells (WT, *vps1*Δ) treated with CSE for different time intervals and were then spotted on YPD agar plates. Each image is representative of ten different fields. Scale bars: 8μm. Results are represented as mean ± SD of data obtained from 10 different fields using paired t test (n.s., not significant; ∗p value < 0.05; ∗∗p value < 0.005; ∗∗∗p value < 0.0005). (a.u. indicates arbitrary unit).

Vps1 plays a vital role for vacuolar fragmentation. To verify that the current study treated *vps1*Δ with CSE and examined the vacuolar morphology and CSE sensitivity. The result showed that after CSE treatment, unlike WT cells, which exhibit fragmentation at 0.5 h, followed by reduction in 1 h and 2 h respectively, *vps1*Δ fails to exhibit vacuolar fragmentation, indicating CSE-induced vacuolar fragmentation is Vps1 dependent (Figure 3D, Left Panel). Quantitative profile also supported this view (Figure 3D, Right Panel).The quantitative result showed that *vps1*Δ exhibited little or no fragment in presence or absence of CSE. Consistent with this, *vps1*Δ cells were found to be more sensitive to CSE compared to wt cells (Figure 3E).

### 2.3. Excessive fragmentation is detrimental to cell

In the previous section, vacuolar fragmentation was shown to be an important component in combating toxic effects of CSE. On the other hand, excessive vacuolar fragmentation has been linked to improper protein sorting across the cell [23]. Therefore, we sought to determine the effects of excessive vacuolar fragmentation on cellular susceptibility to CSE treatment. For this purpose, *ypt7*Δ mutant cells were used, as Ypt7 is known for promoting vacuolar fusion. Consistently, this mutant exhibited excessive fragmentation even without CSE treatment (Figure.4A, Left Panel). Quantitative profile also supported this view (Figure 4A, Right Panel). Thereafter, vacuolar morphology, cell viability, and misfolded proteins were assessed after CSE treatment for different periods. In all instances, the mutant cells exhibited excessive fragmentation, and yes indeed, it was significantly higher compared to wt cells. However, despite this high level of vacuolar fragmentation, mutant cells were found to be more sensitive to CSE than wild-type cells. Again, more misfolded proteins were also found in the mutant cells than wt cells (Figure. 4B, C). These results indicate that there may be a threshold for vacuolar fragmentation below or above which cell survival is affected by CSE-induced oxidative stress.

**Figure 4.**
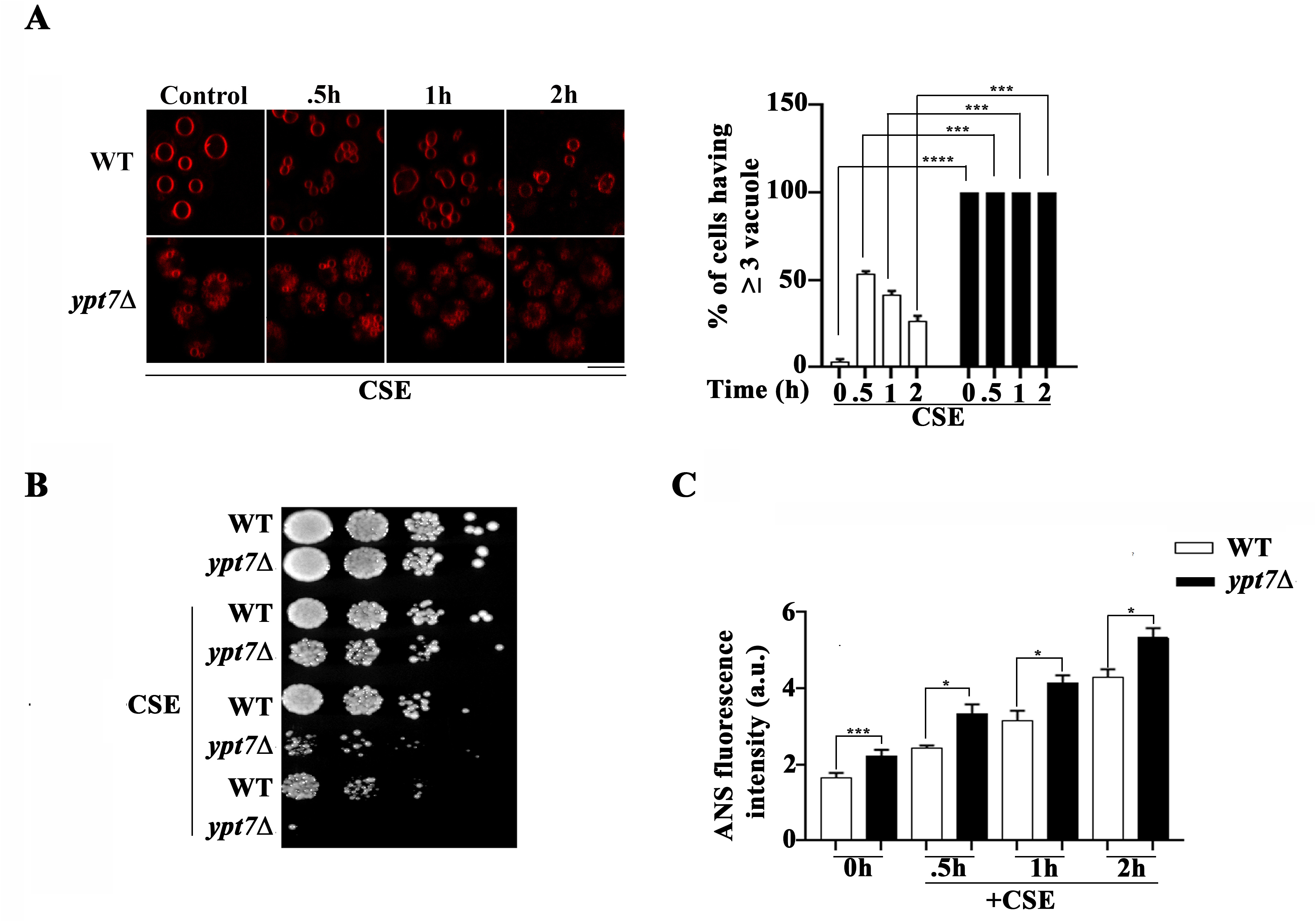
Excessive fragmentation is detrimental for cellular survival during CSE mediated oxidative stress. A) CSE-mediated Vacuolar fragmentation profile for *ypt7*Δ cells. Cells (Wt, *ypt7*Δ) were grown till log phase and were then stained with FM4-64 for 30 mins followed by chasing for 1 h in YPD media. Cells were then treated with CSE for different time periods as indicated and visualized under microscope. Vacuolar morphology of approximately 150 cells was calculated. The bar diagram at the right represent quantitative profile. B) Spot assay. Exponentially growing cells (WT, *ypt7*Δ) treated with CSE for different time intervals and were then spotted on YPD agar plates. C) Misfolded protein Profile. Cells (WT, *ypt7*Δ) were pre grown at 30°C until the early exponential phase and adjusted to an OD_600_ of 0.1. Cells were treated with 15% CSE for different time interval as indicated, followed by washing with PBS and incubated with 30 μM ANS dye for 30 minutes. The level of misfolded protein was determined Spectrofluorimetrically. Each image is representative of ten different fields. Scale bars: 8μm. Results are represented as mean ± SD of data obtained from 10 different fields using paired t test (n.s., not significant; ∗p value < 0.05; ∗∗p value < 0.005; ∗∗∗p value < 0.0005). (a.u. indicates arbitrary unit).

### 2.4. PI 3P and PI (3, 5)P2 play important roles in CSE-mediated vacuolar fragmentation

To verify whether CSE-induced vacuolar fragmentation is PI3P dependent, cells were treated with CSE along with wortmannin, a known PI3 Kinase inhibitor, and as before, vacuolar morphology and cellular survival were examined. The result showed that cells treated with CSE and wortmannin exhibited a significantly reduced fragmentation compared to cells treated with CSE only. Consistent with this, these cells exhibited more sensitivity to CSE compared to cells treated only with CSE (Figure 5A).Interestingly, although treatment of cells with CSE and wortmannin caused reduced fragmentation, these cells displayed prominent vacuolar invagination. It indicates that reduced PI3P levels may not stabilize vacuolar invagination, thereby leading to reduced vacuole fragmentation (Figure 5B).

**Figure 5.**
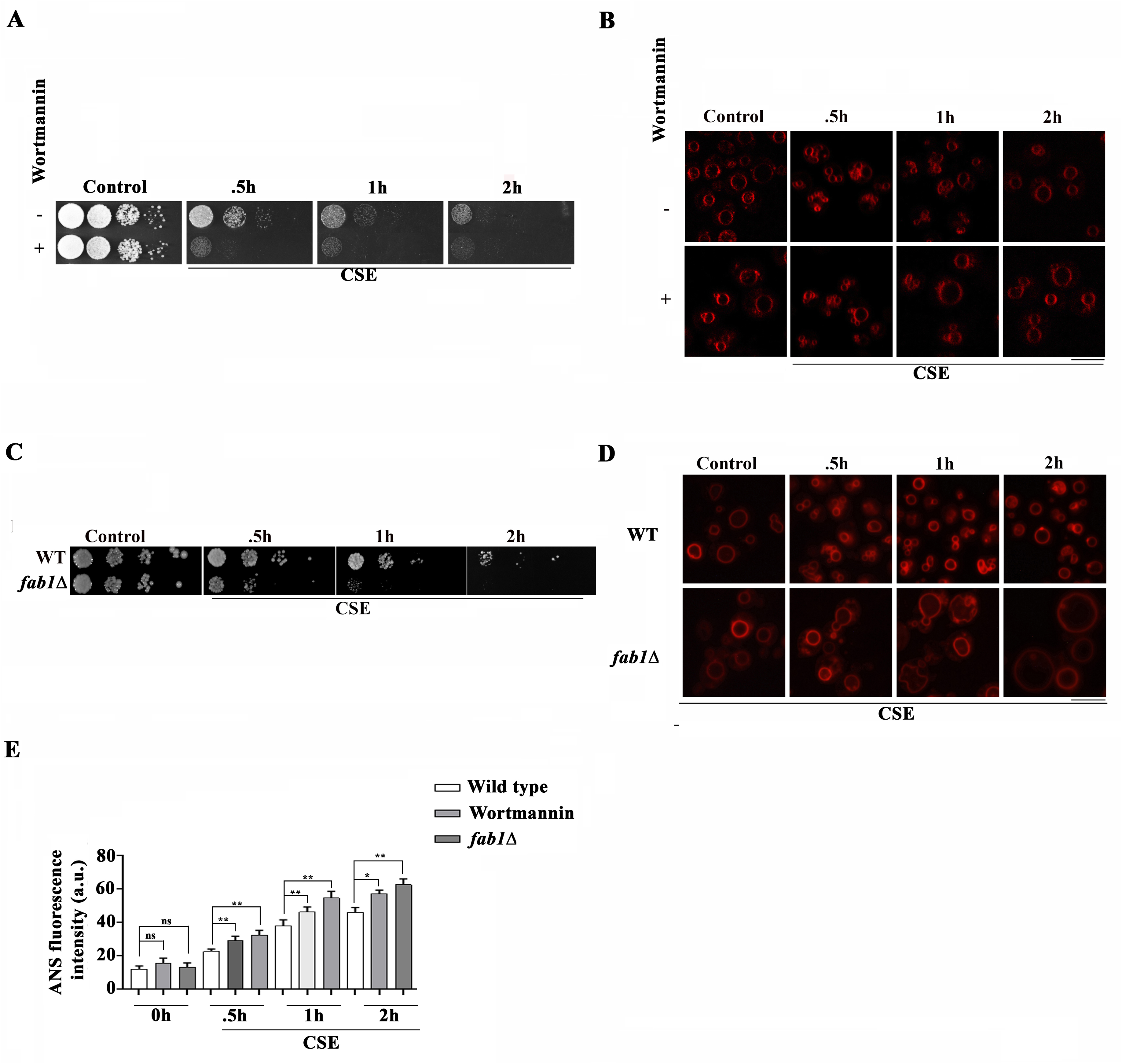
PI3P and PI (3, 5) P2 mediate CSE-induced vacuole fragmentation. **A)** Vacuolar fragmentation profile in wortmannin treated cells. Cells were grown till log phase and were then stained with FM4-64 for 30 mins followed by chasing for 1 h in YPD media. Cells were then treated with CSE in the presence and absence of Wortmannin (100 μM) for different time period as indicated and then visualized under microscope. B) Spot assay. Exponentially growing cells treated with CSE in the presence and absence of Wortmannin for different time intervals and were then spotted on YPD agar plates. C) *fab1*Δ does not undergo vacuolar fragmentation. .Cells (WT, *fab1*Δ) were grown till mid-log phase and were then stained with FM4-64 for 30 mins followed by chasing for 1 h in YPD media. Cells were then treated with CSE for different time interval (.5h, 1 h and 2h) and then visualized under microscope. D) Spot assay. Exponentially growing cells (WT, *fab1*Δ) treated with CSE for different time intervals and were then spotted on YPD agar plates. E) Misfolded protein profile. Cells were pre grown at 30°C until the early exponential phase and adjusted to an OD_600_ of 0.1. Cells were treated with 15% CSE for different time interval as indicated, followed by washing with PBS and incubated with 30 μM ANS dye for 30 minutes. The level of misfolded protein was determined Spectrofluorimetrically. Each image is representative of ten different fields. Scale bars: 8μm. Results are represented as mean ± SD of data obtained from 10 different fields using paired t test (n.s., not significant; ∗p value < 0.05; ∗∗p value < 0.005; ∗∗∗p value < 0.0005).. (a.u. indicates arbitrary unit).

To investigate the role of PI (3, 5) P2 in vacuolar fragmentation, the null mutant of Fab1 (*fab1*Δ), which is a kinase that converts PI3P to PI (3, 5) P2, was examined. The *fab1*Δ cells were treated with CSE and the vacuolar morphology along with their cellular survival were examined. Similar to wortmannin-treated cells, *fab1*Δ cells also exhibited an impaired vacuolar fragmentation and showed greater sensitivity to CSE compared to wt cells (Figure 5C, 5D). In addition to no fragmentation, vacuole size increased progressively with time. The level of misfolded protein load was also measured. As expected, misfolded protein load was found significantly higher in both *fab1*Δ and wortmannin-treated cells after CSE treatment (Figure 5E). Taken together, it can be concluded that PI3P and PI (3, 5)P2 are both important for CSE-mediated vacuolar fragmentation.

### 2.5. CSE-mediated altered lipid homeostasis is important for vacuolar remodelling

The above results indicated potential role of PI3P and PI (3, 5)P2 in vacuolar remodeling upon CSE treatment. However, vacuolar dynamics and their increased surface area are tightly regulated by overall lipid homeostasis. Therefore, the current study investigated the effect of CSE on lipid homeostasis and its contribution to vacuolar fragmentation. To this end, we measured the lipid content of wt cells at different periods (0.5 h, 1 h, and 2 h) after CSE treatment. The result revealed an increase in lipid content at 0.5 h, followed by a decrease at 1 h and 2 h, suggesting that lipid turnover during vacuolar fragmentation is a dynamic process (Figure 6A). To assess the contribution of different cellular lipid pools-phospholipids and stored lipids- we examined the vacuolar morphology of *tgl3*Δ and *plb1*Δ mutants. Tgl3 is a representative of lipases that hydrolyze stored lipids, and Plb1 is a representative of lipases that hydrolyze phospholipids. The *tgl3*Δmutant treated with CSE exhibited a vacuolar fragmentation and growth pattern similar to wt cells. It indicates that the activity of Tgl3 is not required for vacuolar fragmentation. However, it does not rule out the involvement of stored lipids in vacuolar fragmentation, as there exist more isoforms of triglyceride lipase to compensate Tgl3. In contrast, *plb1*Δ mutants failed to undergo vacuolar fragmentation and exhibited higher sensitivity to CSE compared wt cells (Figure 6B, C). This suggests that phospholipids are critical for generating lipids for vacuolar remodeling. It reinforces the hypothesis that vacuolar fragmentation is important for the cellular survival of cells under CSE-mediated stress. Furthermore, given the importance of phosphatidic acid (PA) in membrane curvature and vacuolar fragmentation [24, 25] the level of was also quantified during CSE treatment. At 0.5 h, treatment the level of PA was significantly increased compared to the no-treatment control (Figure 6D). However, beyond 0.5 h, the PA level was progressively decreased mirroring the temporal pattern of vacuolar fragmentation. Based on this observation, it can be said that PA plays an important role in CSE-mediated vacuole fragmentation. Taken together, treatment of cells with CSE alters lipid homeostasis, and this altered pattern plays a major role in mediating CSE-induced vacuolar fragmentation.

**Figure 6.**
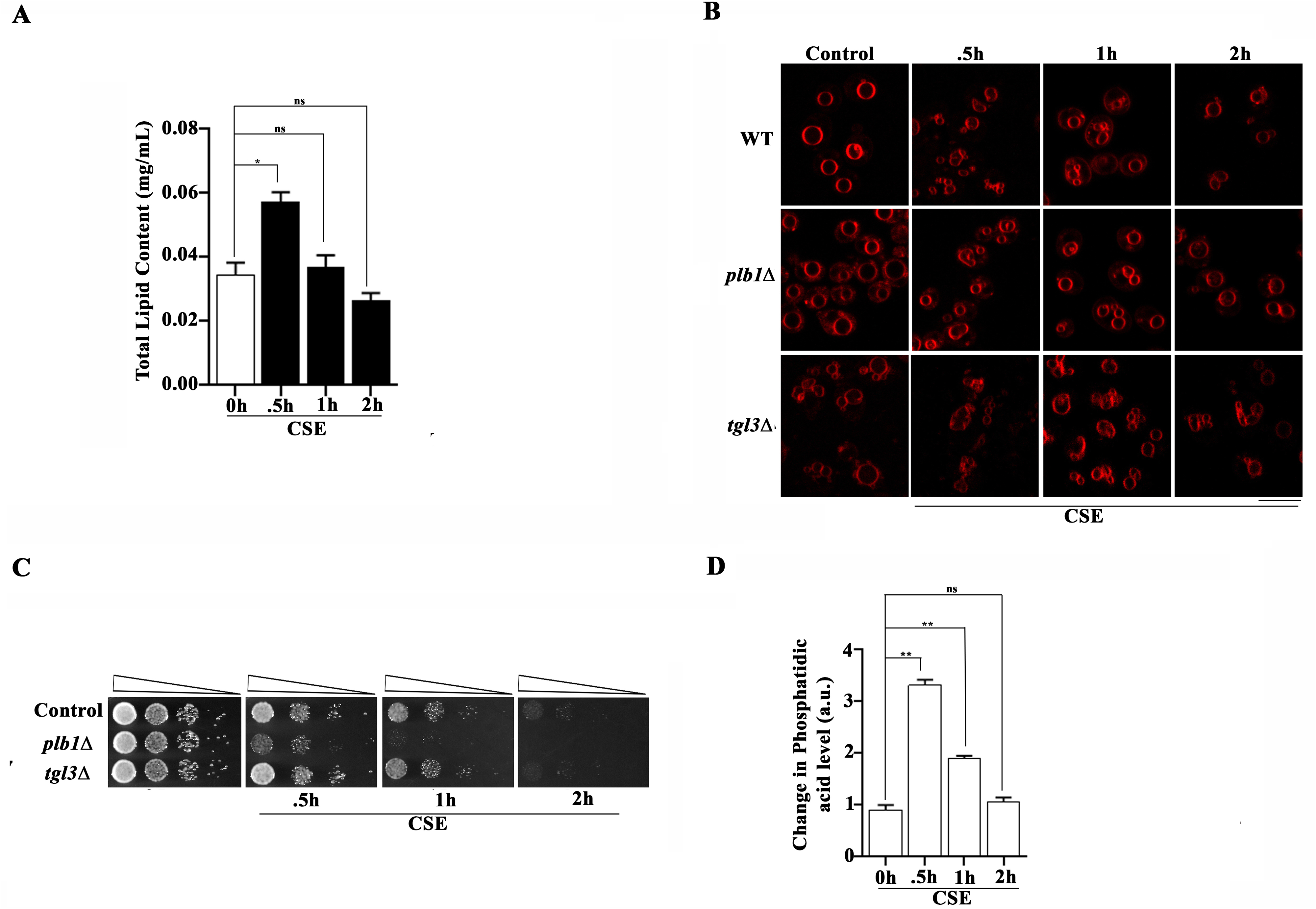
Phospholipase *Plb1* plays a crucial role in CSE induced vacuole fragmentation. A) Cellular lipid profile. Treated and untreated cells pellets were washed twice with Milli-Q water and were dissolve in using chloroform–methanol-water (2.1:1, v/v/v). Total lipid content was measured by gravimetric method after evaporating the chloroform phase. B) Vacuolar fragmentation profile for *plb1*Δ and *tgl3*Δ. Cells (Wt, *plb1*Δ, and *tgl3*Δ) were grown till log phase and were then stained with FM4-64 for 30 mins followed by chasing for 1 h in YPD media. Cells were then treated with CSE for different time period as indicated and then visualized under microscope. C) Spot assay. Exponentially growing cells treated with CSE for different time intervals and were then spotted on YPD agar plates. D) Phosphatidic acid profile. Treated and untreated cells pellets were washed twice with miliQ water and were dissolved in Methanol: Diethyl ether: Saturated salt (1:1:1). The Diethyl ether fraction was then collected and run on HPLC. Each image is representative of ten different fields. Scale bars: 8μm. Results are represented as mean ± SD of data obtained from 10 different fields using paired t test (n.s., not significant; ∗p value < 0.05; ∗∗p value < 0.005; ∗∗∗p value < 0.0005). (a.u. indicates arbitrary unit).

### 2.6. Mitochondrial activity is important for vacuolar remodeling

The effect of mitochondrial respiration on vacuole fragmentation was examined as it provides energy necessary for vesicle transport [26, 27]. To test this, cells were treated with mitochondrial respiration inhibitor oligomycin along with CSE and were looked for their vacuolar morphology as well as sensitivity towards CSE. Oligomycin treated cells exhibited both excessive sensitivity and a reduced vacuolar fragmentation compared to cells that are treated only with CSE at all time points with the difference becoming more pronounced after 1 h (Figure 7A,B). It suggests that reduced availability of energy inside the cell lowers vacuolar fragmentation.

**Figure 7.**
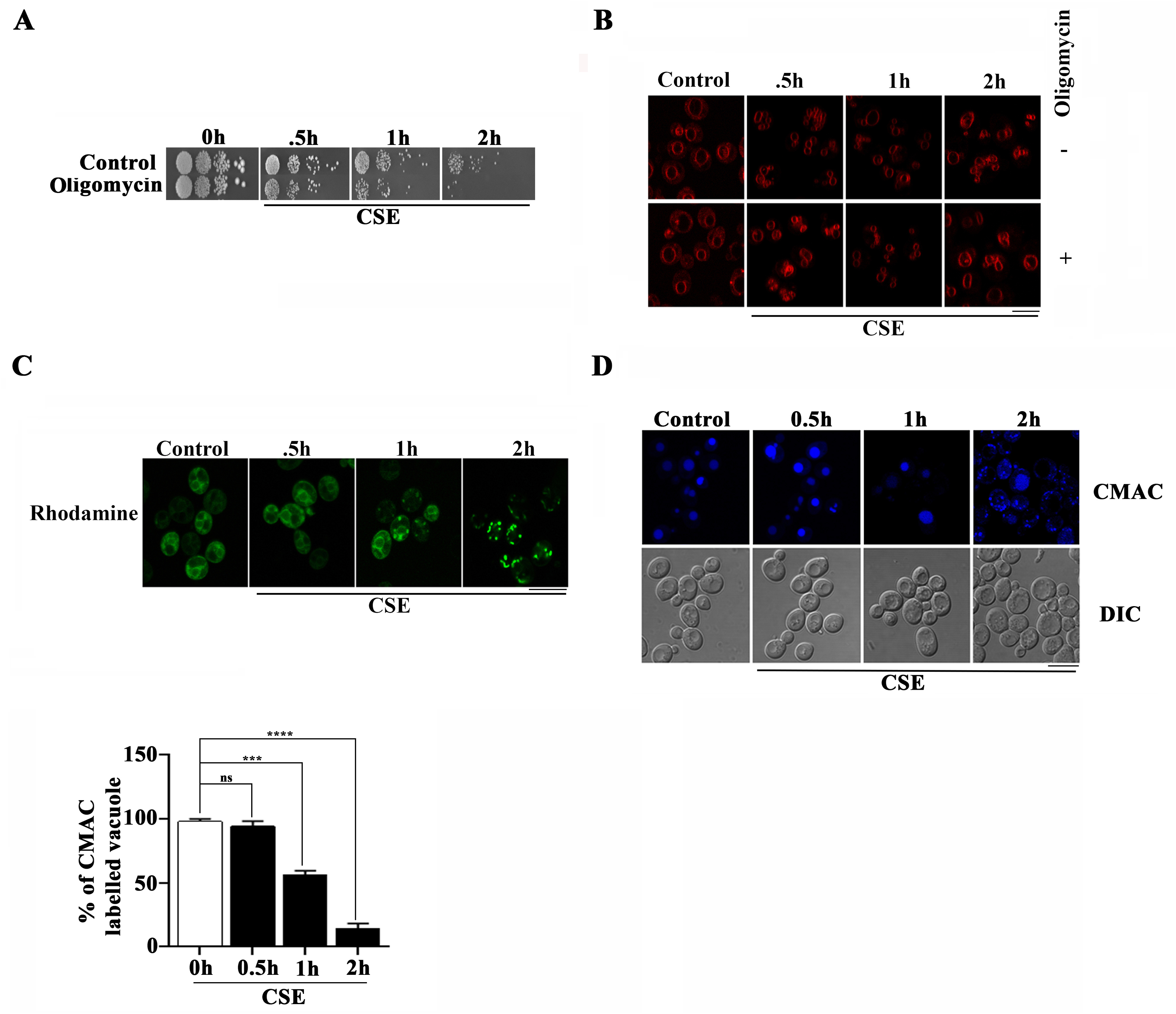
Functional mitochondria are required for CSE-mediated vacuolar fragmentation. A) Spot assay. Cells were grown till mid log phase and treated with 15% CSE for different time period as indicated in the presence and absence of mitochondrial inhibitor. Cell was then spotted for growth. B) Vacuolar fragmentation profile in presence of oligomycin. Cells were grown till mid-log phase and were then stained with FM4-64 for 30 mins followed by chasing for 1 h in YPD media. Cells were then treated with CSE for different time period as indicated in the presence and absence of oligomycin (10 μm) and then visualized under microscope. C) Rhodamine123 staining profile. Cells were grown till mid log phase and treated with CSE for different time period as indicated and were then stained with Rhodamine123 and were observed under microscope. D) CMAC staining profile. Cells were grown till mid log phase and treated with CSE for different time period as indicated and were then stained with CMAC dye and were observed under microscope. The bar diagram at the right side represents the quantitative profile. Each image is representative of ten different fields. Scale bars: 8μm. Results are represented as mean ± SD of data obtained from 10 different fields using paired t test (n.s., not significant; ∗p value < 0.05; ∗∗p value < 0.005; ∗∗∗p value < 0.0005).

If this is the case, then the observed reduced fragmentation after 0.5 h of treatment may also arise from impaired mitochondrial activity. To investigate this, the mitochondrial membrane potential was assessed by staining CSE treated cells with Rhodamine 123 dye, a dye taken up by mitochondria due to its negative membrane potential producing green fluorescence. Green fluorescence was observed for all the time points but its intensity and the pattern varied.

Untreated cells and those treated for 0.5 h displayed a filamentous mitochondrial morphology. At 1 h, a mixed phenotype appeared with some cells resembling untreated controls and a substantial population exhibited punctuated green structure. At 2 h nearly all cells exhibited punctated mitochondria. The formation of punctated structures, rather than a filamentous pattern, suggests a morphological alteration of mitochondria in CSE-treated cells, potentially leading to reduced energy efficiency (Figure 7C). To verify this functional decline, we examined V-ATPases assembly by examining vacuolar pH. An appropriate V-Atpases assembly is required for both vacuolar fragmentation and the maintenance of pH and it is energy dependent. A defect in mitochondrial respiration will therefore impair V-Atpases assembly that in turn perturbs vacuolar pH homeostasis. Vacuolar pH was monitored using CMAC dye, which fluoresces brightly at the physiological pH of the vacuole.CSE treated cells for 1 h or more showed impaired CMAC staining indicating altered vacuolar pH. This finding supports the idea that beyond 0.5 h of CSE treatment both V-Atpases activity and mitochondrial function are compromised (Figure. 7D). . Taken together, these results indicate that mitochondrial respiration declines after 0.5 h of CSE treatment, contributing to reduced vacuolar fragmentation.

## 3. Discussion

The present study elucidates the impact of CSE on vacuolar physiology and establishes finely tuned vacuolar fragmentation as a critical adaptive mechanism to combat cigarette smoke induced oxidative stress, [15, 28]. We demonstrate that this process is an active, advantageous event that is necessary for survival and proteostasis. It is controlled by specific lipid signaling and membrane remodeling. This mechanism is energy-intensive and transient as it is disintegrated under extended stress because of mitochondrial malfunction. The study provides a model that integrates organelle dynamics, lipid remodeling, and energy balance with stress adaptation.

Why would a stressed cell expend energy breaking apart a functioning organelle? Our research demonstrates that vacuolar fission optimizes the vacuole for microautophagy by raising the surface area, which improves proteostasis. We suggest that oxidative stress causes increased vacuolar membrane tension. This increased membrane tension might promote mechanical scission, leading to vacuolar fragmentation, ensuing a larger vacuole surface area. This shape is essential because it optimizes the interface for direct microautophagic engulfment of cytoplasmic cargo, including aggregates of misfolded proteins, when canonical protein clearance pathways are overloaded [15]. Therefore, it is crucial to understand how oxidative stress is converted into a targeted membrane remodeling event. To this end, we find that a specific lipid signaling holds the key. The early increase in phosphatidic acid (PA) and requirement of PLB1 activity suggest that lipid hydrolysis provides the substrates that cause curvature in fission. Phosphatidic acid (PA) is a crucial lipid modulator of membrane dynamics, including vacuolar fission [24,25].PA provides negative membrane curvature, which is a biophysical need for membrane constriction and fission processes [24,25].PA acts as a spatial cue to attract fission machinery by preferentially accumulating at highly curved membrane domains and subsequently helps them get activated. Furthermore, PA serves as a major metabolic axis because it is a precursor that produces lysophosphatidic acid (LPA) and diacylglycerol (DAG), both of which have an impact on membrane remodeling. In response to exposure to CSE, PA may accumulate at vacuolar membranes as its level was increased. This could be a synergistic process in which PA might work in tandem with PI (3,5)P_₂_ and oxidative stress signaling to cause vacuolar fragmentation.

One of the key findings of our study was the identification of PI (3, 5) P_2_ as an essential mediator of vacuolar fragmentation in response to CSE exposure. PI3P and PI (3, 5) P_2_ are involved in the regulation of vacuolar membrane dynamics and are crucial for both vacuolar fission and fusion [29, 30, 31, 32]. PI (3, 5) P2, which is produced from PI3P by Fab1 kinase, plays a vital role in the fragmentation process by allowing three crucial processes. First, it recruits curvature-sensing proteins like Atg18 to stabilize membrane tension. Secondly, it targets the GTPase Vps1 to membrane neck regions by interacting with the PH domain of VPS1and thereby promoting the vacuolar scission. Third, phospholipase PLB1 (Figure 6F) is activated by PI (3, 5) P_₂_, releasing fatty acids required for the formation of phosphatidic acid (PA). A feed-forward loop is created by this action, which facilitates more membrane expansion. These results clarify why scission is not permitted by wortmannin treatment, which prevents PI (3, 5) P_₂_. Our finding is consistent with a previous report showing vacuolar fragmentation caused by ROS accumulation from excess iron, zinc, or copper, and it becomes exacerbated by the loss of mitochondrial SOD2 [33]). Interestingly, treatment with rapamycin inhibits metal-induced fragmentation. Since rapamycin is a known inhibitor of TORC1 and thus all these things suggest that metal homeostasis, oxidative stress, and TORC1-dependent signaling are all linked to vacuolar morphology in fungi. Notably, cells were sensitized to CSE by both excessive fusion (via YPT7 overexpression) and excessive fragmentation (by *ypt7*Δ), indicating that the ideal vacuolar dynamics is necessary for better survival (Figures. 3–4). Hyper-fragmentation can interfere with endosomal trafficking, cargo sorting and also essential vacuolar functions such as major storage compartment, osmoregulation and ion homeostasis [22], while excessive fusion reduces vacuolar surface area and prevents microautophagic interaction. The inability of *vps1*Δ cells (Figure. 3D-E) to perform fission [29] due to their hypersensitivity highlights the non-redundant function of fission machinery under oxidative stress. However, it is unclear, how cells recognize and preserve fission-fusion equilibrium. Despite the initial advantages, vacuolar fragmentation is temporary. By two hours of CSE treatment, fragmentation decreases despite ongoing stress, which is consistent with lower mitochondrial function (Figure. 7C-D). Vesicular trafficking, fission machinery activity, and V-ATPases assembly and acidification (Figure. 7E) are all ATP-dependent processes that become dysfunctional by this energy collapse [26, 27]. As a result, misfolded proteins build up, and proteostasis breaks down and finally cells undergo death. This phenotype is exacerbated by oligomycin (Figure. 7A-B), confirming mitochondria as the limiting component. As can be seen, the phenotype of *sod1*Δ yeast under oxidative stress is similar to the vacuolar fragmentation brought on by exposure to cigarette smoke extract (CSE) [34]. Elevated amounts of reactive oxygen species (ROS) and labile intracellular iron combine to harm vacuolar membranes in *sod1*Δ mutants. As a Fenton catalyst, iron transforms H_2_O_₂_ into very reactive hydroxyl radicals that damage membranes and proteins. Similarly, by redistributing or redox-activating vacuolar Fe^2+^ pools, CSE may encourage localized oxidative damage. Vacuoles are important locations for storing iron; therefore their fragmentation may act as a compensatory strategy to improve iron sequestration by boosting surface area and recruiting transporters.

In conclusion, vacuolar fragmentation emerges as a survival mechanism for cell exposed to stress. However, its failure under prolonged stress highlights an inherent trade-off: While it provides a quick adaptation, it is also energy-intensive and prone to collapse. In this study, we demonstrate that PLB1 and PI (3, 5) P2 are key players in vacuolar membrane remodeling during CSE-induced oxidative stress. Enhancing PLB1 and PI (3, 5) P2 activity may improve cell survival by facilitating the clearance of misfolded proteins that accumulate in CS-induced oxidative stress. Thus, PLB1 and PI (3,5) P2 represent potential therapeutic targets in diseases where misfolded protein clearance is essential. Importantly, the translational significance of this yeast model is underscored by the conservation of the PI3K/Fab1 pathway and mTOR-sensitive vacuolar remodeling in mammalian cells.

## 4. Materials and Methods

### 4.1. Preparation of cigarette smoke extracts (CSE)

CSE was prepared using a commercially available filter-tipped cigarette with a tar content of 15 mg and nicotine content of 1 mg manufactured by the Indian Tobacco Company Limited (ITC Ltd.) as described previously. The smoke from a cigarette was bubbled through 1 ml of 50 mM PBS, pH 7.4, till the cigarette was consumed completely. The pH of this aqueous CSE was adjusted to 7.4 using NaOH. The OD_0.6-0.8_ was considered 100%, which is diluted as per requirements.

### 4.2. Yeast strain, media and CSE treatment

*Saccharomyces cerevisiae* (*S. cerevisiae*) strain BY4741 (*MAT*a *his3*Δ1 *leu2*Δ0 *met15*Δ0 *ura3*Δ0) was used in this study. All the null mutants used in this study have BY4741 genetic background and were obtained from Prof. Maya Schuldiner (Weizmann Institute of Science, Israel). HSP104-GFP also have BY4741 genetic background and this was a kind gift from Prof. Markus Tamas (University of Gothenburg, Sweden). Yeast cells were grown in synthetic drop-out medium that contains 2% glucose and 0.1% amino acid mixture (containing arginine – 0.9 g, methionine – 0.9 g, tyrosine – 1.35 g, isoleucine – 1.35 g, phenylalanine – 1.35 g, aspartic acid – 4.5 g, glutamic acid – 4.5 g, valine – 4.5 g, threonine – 4.5 g and serine – 4.5 g) along with 0.005% lysine. As for the CSE treatment, based on the previous reports [16] exponentially growing *S. cerevisiae* cells were treated with 15% CSE for different periods

### 4.3. Spot assay

Exponentially growing *S. cerevisiae* cells (10^6^/mL) were treated with 15% CSE for 0.5 h, 1 h and 2□h in the presence or absence of additional reagents as required for the specific experiments. Cells were centrifuged, resuspended in fresh medium and serially diluted. Thereafter an aliquot from the serially diluted samples were spotted on the appropriate agar plates. Plates were incubated at 30□°C for 48□h.

### 4.4. Colony forming unit (CFU) assay

Exponentially growing *S. cerevisiae* cells (10^6^/mL) were treated for with 15% CSE for different periods, with or without additional reagents as required for the specific experiments. Cells were harvested, resuspended and serially diluted in fresh medium and thereafter 100μL was spread on the agar plates. The plates were then incubated at 30□°C for 48 h. Colonies were counted and CFU/mL was calculated.

### 4.5. Determination of protein misfolding by ANS dye

Misfolded protein level was measured spectrophotometrically as described by [17]. Briefly, cells were grown up to mid log phase and the O.D value was set to 0.1. Cells were treated with 15% CSE for 2 h. Then, cell lysate were prepared with lysis buffer (Tris-HCl-50mM, EDTA-5mM, NaCl-250mM, NP-40-0.1%, DTT-1.5mM, PMSF-1mM, protease inhibitor cocktail-1:1000 ratio). Further, protein concentration was measured by Lowry method [18]. An aliquot containing 10μg of protein was taken from each lysate for determination of protein misfolding by ANS dye. ANS dye was added to each lysate at a final concentration of 30 µM and incubated in the dark at room temperature for 30 mins. Fluorescence was measured in a spectrofluorimeter (Hitachi F7000, Japan) (Excitation: 420nm, Emission: 450-550nm).

### 4.6. FM4-64, CMAC and Rhodamine Staining

For FM4-64 staining, cells were stained with FM4-64 before CSE treatment. Cells were grown to logarithmic phase and centrifuged. The pellet was resuspended in 100 μL of YPD (Yeast extract 1%, Peptone 2%, Dextrose 2%), and 0.5 μl FM4-64 solution (30 µM) was added to it. It was kept in the dark under shaking conditions for 45 mins. Cells were then spun down, washed and resuspended in 850 μL YPD medium and again kept for 1 h. Thereafter, to it 150 μL of CSE was added to make the final concentration 15% and incubated for different periods as indicated in the figures, followed by washing with YCM media, and were visualized under a fluorescence microscope (Olympus IX 71, Japan).

For CMAC staining, exponentially growing cells were treated with CSE as mentioned in the previous section, washed and then incubated with CMAC at a final concentration of 5 μM at room temperature in the dark for 30 minutes. Cells were then washed and visualized under a confocal laser scanning microscope (Leica Stellaris 5, Leica Mikrosysteme Vertrieb GmbH, Wetzlar, Germany).

For Rhodamine staining, cells were treated with CSE as before, washed and then incubated with Rhodamine 123 (final concentration 5 mg/mL) for 20 min. Cells were centrifuged, washed, resuspended in PBS and then visualized under a confocal laser scanning microscope (Leica Stellaris 5, Leica MikrosystemeVertrieb GmbH, Wetzlar, Germany).

### 4.7. Fluorescence and Confocal microscopy

To visualize GFP-tagged proteins, FM4-64, CMAC and Rhodamine 123 dye stained cells either a regular fluorescence microscope (Olympus IX 81, Japan) or a confocal laser scanning microscope (Leica Stellaris 5, Leica MikrosystemeVertrieb GmbH, Wetzlar, Germany) was used as described in the results section. For GFP-tagged proteins and Rhodamine 123 dye stained cells, the FITC filter was set with an excitation wavelength of 395 nm and an emission wavelength of 510 nm. For FM4-64-stained cells, the TRITC filter was set with 515 nm for excitation and 640 nm for emission.

### 4.8. Extraction of total lipid

Total lipids were extracted following the phase separation method of Bligh and Dyer [19] dissolved using chloroform–methanol-water mixture (2.1:1, v/v/v). Total lipid content was estimated by the gravimetric method after evaporating the chloroform phase.

### 4.9. Estimation of Phosphatidic Acid (PA)–

Phospholipids were extracted from the cell lysate using methanol -diethyl ether -saturated NaCl mixture (1:1:1, v/v/v). After phase separation, the diethyl ether fraction was collected. An aliquot of it was loaded on HPLC column. HPLC machine (Waters, Singapore) consists of a 1525 binary HPLC pump, and a 2475 multi-wavelength fluorescence detector. The reverse phase C8 (4.6 × 250 mm; Waters) analytical column was used. The mobile phases were a mixture of methanol, acetonitrile and water (90.5: 2.5,7, v/v/v). A pure PA solution was used as standard.

## 5. Statistical analysis

All experiments were performed at least three times independently, at least 150 cells were assessed from each, and the results were expressed as mean value ± standard deviation (SD). To find the statistically significant differences, paired t test was performed using GraphPad Prism software [Version 8.0.2(263)]. Differences were considered statistically significant at p<0.05.

## Author contributions

AS designed experiments, performed all the experiments, analysed the results and wrote the primary manuscript. SS participated in designing experiments and analysed the results. AKS conceived the idea, designed experiments, analysed results, and revised the manuscript.

## Acknowledgement

The authors would like to acknowledge Prof. Maya Schuldiner (Weizmann Institute of Science, Israel) and Prof. Markus Tamas (University of Gothenburg, Sweden) for providing *Saccharomyces cerevisiae* strains used in this study. The authors would also like to acknowledge the Council of Scientific and Industrial Research (CSIR), Govt of India, for providing financial support to AS [09/028(1111)/2019-EMR-1]. The authors would also like to acknowledge the Bose Institute Central Instrumentation Facility and IPLS (University of Calcutta) Instrumentation Facility.

## Ethics Statement

The authors have nothing to report.

## Data availability

All the data are included in the manuscript.

